# Identification of small-molecule enhancers of circadian rhythm amplitude in central and peripheral clocks

**DOI:** 10.64898/2026.07.23.740283

**Authors:** Maitreyi S. Joshi, Felix P. Albert, Johanna H. Meijer, Stephan H. Michel, Paul P. Geurink

**Author notes:** Correspondence: **Maitreyi S. Joshi, Email:**.

## Abstract

The circadian clock regulates daily rhythms in metabolism, behavior, and physiological homeostasis. Damped or disrupted circadian rhythms are linked to physiological malfunctions, including neurodegeneration, metabolic syndrome, and cancers. Enhancing circadian rhythm amplitude can strengthen the processes regulated through the circadian system and offer a potential therapeutic approach. Here, we present eight small-molecule enhancers of circadian amplitude with previously unrecognized roles in circadian regulation. By using a *Bmal1* transcription reporter system in NIH3T3 fibroblasts, we screened a drug repurposing library comprising 5,631 diverse compounds. Through which eight molecules were identified to increase circadian amplitude across multiple cycles, in a dose-dependent manner, without significantly perturbing period or phase. We evaluated the physiological potential of these compounds in ex vivo mouse tissues expressing PER2 luminescence reporter. A set of compounds tested in organotypic liver or suprachiasmatic nucleus slices enhanced the circadian amplitude in respective tissues, showing their suitability for peripheral as well as central clocks. Single-cell imaging of suprachiasmatic nucleus slices revealed that amplitude enhancement results from increased expression of clock proteins in individual cells rather than from intercellular synchronization. Together, these compounds and their targets present a new array of circadian amplitude modulators that may present opportunities to pharmacologically enhance clock robustness in physiology and disease.

**Significance Statement:** Through a network of molecular interactions, the circadian clock coordinates rhythms of biochemical and behavioural processes, right from the level of a single cell to organs. Therefore, maintaining a strong circadian clock, through its high-amplitude circadian rhythm, is crucial. However, few pharmacological approaches and targets are known to enhance circadian amplitude. This study presents eight previously uncharacterized amplitude enhancer molecules by conducting a chemical screen on cultured fibroblasts. Identified molecules also induced amplitude enhancement in mouse-derived liver and brain-SCN slices, showing their robust functionality across tissues. In brain-SCN slices, the amplitude enhancement was found to occur on the single-cell level without affecting the synchronization between cells.

## Introduction

The circadian clock is an intrinsic time-keeping system that orchestrates the temporal organization of a wide range of physiological processes, including sleep-wake cycles, hormonal signaling, metabolism, and cellular proliferation (1, 2). Disruption of these rhythms is associated with diverse pathological conditions, highlighting the importance of robust circadian regulation. Relevant features of circadian rhythms are period, phase, and amplitude. Here, period is defined as the time span of one oscillatory cycle (∼24 hrs), phase is described in the context of relative temporal alignment (or synchronization) between multiple rhythms, and amplitude is the difference between an adjacent peak and trough in the oscillation (Fig. 1A). Amplitude signifies the strength of oscillations of clock protein expressions and therfore how strongly it will connect with various clock coupled biochemical processes in the body (e.g., cell cycle) (3). This is particularly important as the circadian clock regulates the oscillating expression of several receptors, enzymes, and metabolites, often termed clock-controlled proteins. As the circadian clock tightly regulates metabolic gene expression, damped circadian rhythms have been shown to cause metabolic disturbances such as hyperlipidaemia and hyperglycaemia (4). In the elderly, low-amplitude rhythms have been linked to susceptibility to dementia and loss of temporal coherence in the metabolome, signifying amplitude as a measure of the circadian rhythm robustness and overall physiological resilience (5, 6). On the other hand, high-amplitude organ-level circadian clocks are often associated with improved functionality, as demonstrated in liver metabolism (7). Therefore, identifying novel pharmaceutical means to enhance the amplitude of the circadian rhythm will present opportunities to improve clock-controlled physiological outcomes.

**Figure 1:**
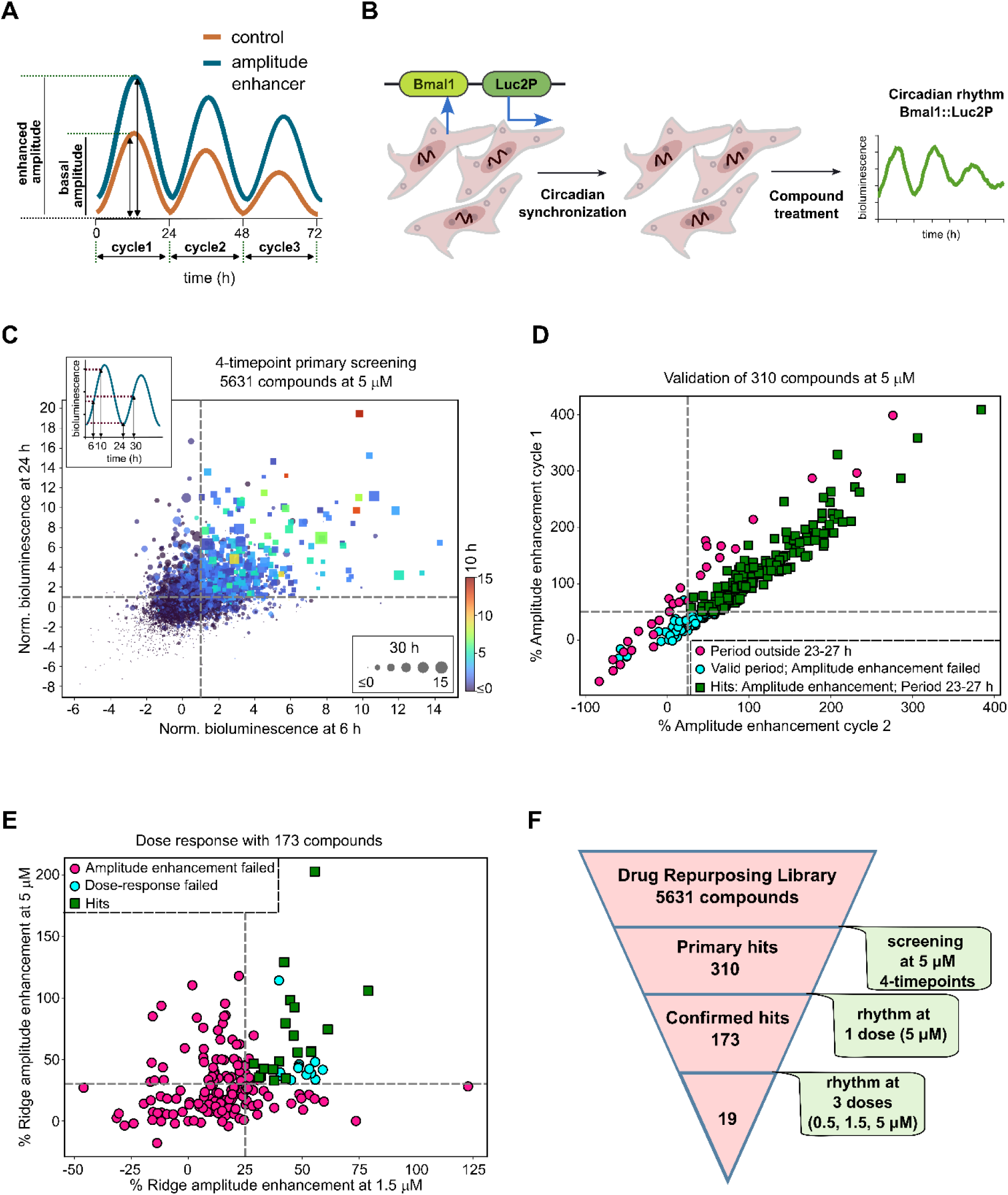
Cell-based high-throughput screening for identifying circadian rhythm amplitude enhancer molecules. (A) Illustration of the change in molecular circadian rhythm upon administration of an amplitude-enhancing small-molecule. (B) Recording of circadian rhythm in cells with bioluminescence readout based on NIH3T3 mouse fibroblasts expressing Bmal1::Luc2P. (C) Four-dimensional scatterplot representing results of the primary screen of 5,631 compounds at 5 μM. Bmal1::Luc2P bioluminescence from phase-synchronized NIH3T3 cells was recorded at four time points (6, 10, 24, and 30 h, indicated in the top left inset) after treatment with compounds. Each circle/rectangle represents luminescence values normalized to the DMSO average and standard deviation at the respective time points. Normalized luminescence at 6 and 24 h is represented on X- and Y-axes, respectively, 10 h values are represented across the indicated color spectrum, and 30 h with marker sizes. A compound is termed a hit (represented with rectangles) if its normalized bioluminescence value is 1-time above DMSO-standard deviation at 6, 24, 30 h and 1.5-time above DMSO-standard deviation at 10 h; or else it is a non-hit (circles). (D) Scatter plot showing validation of 310 hit compounds from (C) by applying them to cells at 5 μM and recording Bmal1::Luc2P circadian rhythm for 3.5 days. Compounds are hits (green rectangles) if they show amplitude enhancement above DMSO in cycle1 and cycle2, as well as across the whole rhythm, and their periods lie within 23-27 h. Compounds are non-hits if they fail the amplitude (cyan circles) or period criteria (magenta circles). (E) Scatter plot showing dose response of 173 hit compounds from (D) by applying them to cells at 0.5, 1.5 and 5 μM and recording Bmal1::Luc2P circadian rhythm for 3.5 days. Compounds are hits (green rectangles) if they qualify amplitude enhancement criteria in (D) at 1.5 and 5 μM, and they show a dose-dependent amplitude enhancement. Compounds are non-hits if they fail the amplitude (magenta circles) or dose-response criteria (cyan circles). (F) Schematic overview of the sequential high-throughput screening and validation workflow with identified hits indicated for each step.

The circadian system is roughly divided into the suprachiasmatic nucleus (SCN) clock in the brain, commonly referred to as the central clock, and non-SCN clocks, referred to as peripheral clocks (8). At the level of single cells, both central and peripheral clocks make use of a core negative feedback loop formed between a set of genes: *Bmal1*, *Clock*, *Per1/2,* and *Cry1/2,* whose transcription-translation-repression gives rise to circadian oscillations of their protein expression (known as the transcription-translation feedback loop (TTFL); SI Appendix, Fig. S1). The transcription events in the primary feedback loop are coupled to a secondary feedback loop formed by nuclear receptors like REV-ERBs and RORγ, which adds a further layer of regulation to molecular circadian rhythms (9). Considering the contribution of various proteins and interactions towards the regulation of rhythm parameters (period, phase, and amplitude), a few small-molecule clock modulators have been identified that act on primary or secondary loop elements. This includes stabilizers of CRY (e.g., KL001) to achieve period lengthening, as well as agonists of RORα/γ (e.g., nobiletin) or inhibitors of CLOCK-BMAL1 interaction (e.g., CLK8) to enhance the rhythm amplitude (10–12). Moreover, post-translational modifications (e.g., phosphorylation through kinases like CK1) and other intracellular oscillators (e.g., NAD^+^ biosynthesis, cAMP signaling) affect the core clock elements and regulate their interactions, activities, and localization, which is shown to modulate the circadian rhythm (13–15). Therefore, it is necessary to widen our perspective from TTFL and search for novel targets outside the core-clock network.

While investigating circadian rhythm modulation, chronobiological studies often focus on period and phase because of the ease of their direct measurement. This has also led to years of investigation into the identification of genetic factors, protein targets, and post-translational processes that affect the circadian period and phase. For example, phosphorylation of PER and CRY proteins through Casein kinase activities like CK1 or CK2 is found to be crucial for the precise modulation of circadian period (13, 16). Knowledge of strong nodes like these kinases has also led to target-directed chemical screens to identify and improve small molecules to achieve period modulation in the desired direction. In contrast, mechanisms regulating circadian amplitude have received less attention, making it difficult to target specific proteins and activity sites to modulate circadian amplitude. This opens a window of opportunity to explore novel targets and to learn about biochemical nodes outside the core clock network that can be exploited to achieve circadian amplitude modulation. An efficient way to identify novel targets is through high-throughput screening with small-molecule libraries with diverse targets. This method was previously used to identify amplitude-enhancing molecules like CEM3, nobiletin, and ISX9 (17, 18).

Here, we aimed to identify novel compounds and protein targets that could enhance the amplitude of the molecular circadian rhythm without significantly affecting its period and phase. To find useful targets from diverse categories of compounds, we conducted a target-agnostic high-throughput screen using a drug repurposing library. We adapted live-cell circadian reporter assays to a high-throughput environment to monitor the effect of screened molecules on the circadian oscillations of core clock gene *Bmal1*. This resulted in the identification of 13 small molecules with diverse structural motifs as molecular circadian rhythm amplitude enhancers. Amongst these, eight molecules or their corresponding published targets were previously unknown to affect circadian amplitude. The amplitude-enhancing potential of these compounds was validated on mouse liver- and brain SCN-organotypic slices. Using this approach, we identified eight compounds that effectively enhanced circadian amplitude not only in cultured fibroblasts but also in liver (peripheral) tissues, with six of these compounds additionally showing applicability for amplitude enhancement in the SCN (central) clock.

## Results

### High-throughput screen for modulators of circadian rhythm amplitude

We adapted a high-throughput assay to record real-time circadian rhythm in NIH3T3 mouse fibroblasts (10, 16–19). These cells were chosen for their stable genetic background and prior characterization of the molecular circadian rhythm. To monitor the rhythm modulatory effect of compounds on a core clock component-BMAL1, we transfected a luciferase reporter plasmid (Bmal1::Luc2P) under the control of the Bmal1 promoter (Fig. 1B). This system serves as a parallel readout for the changes in the transcription of Bmal1. With this, we screened a drug-repurposing library comprising 5,631 off-patent compounds with a diverse range of targets. To screen such a large number of compounds efficiently, we first measured Bmal1::Luc2P bioluminescence at four time points across the circadian profile upon sustained compound treatment (5 µM) instead of recording the high-resolution circadian rhythm profile at this stage (18). In this primary screen, we recorded the bioluminescence at 6 hr, 10 hr, 24 hr, and 30 hr post-compound administration and formulated a screening criterion based on the enhancement in bioluminescence above vehicle (DMSO) treatment (Fig. 1C). This approach allowed us to identify compounds that have a reproducible effect on circadian rhythm amplitude. To identify amplitude enhancers, the bioluminescence signal at each time point was normalized to the average change induced by DMSO, and compounds were termed as potential amplitude enhancers if they increased the bioluminescence 1-times above the standard deviation of DMSO at 6 hr, 24 hr, and 30 hr, as well as 1.5-times above the standard deviation of DMSO at 10 hrs. As the signal-to-noise ratio was higher near the rhythm peak (∼10 hrs) than at time points farther from the peak, a more stringent screening criterion was applied at the 10-hour time point. This resulted in the selection of 310 hit compounds with possible amplitude-enhancing properties.

The primary screen does not filter out false positives that might be identified as amplitude enhancers because of the buildup of BMAL1-bioluminescence, inducing a lack of rhythicity or large deviations in circadian period. Therefore, we applied the 310 hit compounds to cells at a concentration of 5 µM and recorded the complete waveform of the Bmal1::Luc2P rhythm for 3.5 days (sample interval 10 min; Fig. 1D). Each recorded bioluminescence timeseries post-compound treatment was analyzed with a continuous wavelet transform to extract rhythm parameters (see methods). By imposing criteria to select for compounds that enhance the amplitude across the circadian cycles (>50% amplitude enhancement in cycle1 and >25% in cycle2), maintain rhythmicity (peak detection), and do not induce large period modulation (period between 23-27 hrs), we picked 173 compounds for the next round. These compounds were then applied to cells at three doses: 0.5, 1.5, and 5 µM and qualified as hits if they showed a dose-dependent effect on amplitude in addition to being consistent amplitude enhancers throughout the circadian cycles (Fig 1E). With this, 19 compounds met all criteria, 13 failed to show dose-response upon increasing concentration, and 141 compounds failed the amplitude-enhancement criteria at either 1.5 or 5 µM. Together, we obtained 19 final hits across diverse chemical scaffolds as amplitude enhancers (Fig. 1F, SI Appendix).

### Eight novel amplitude enhancers were identified in NIH3T3 fibroblasts

To rule out the possibility of contamination of compounds in the library plates, we verified the amplitude-enhancing effect of hit compounds using freshly ordered commercial stocks (Figs. 2 and SI Appendix, Fig. S2A). Out of the 19 obtained hits, 6 compounds-Isrib transisomer, Ruxolitinib, PD53035, ON123300, PQ401 and Bay K8644 failed to show amplitude enhancement (Fig. 2A-F). Whereas, 13 compounds emerged as true-positives by inducing more than 25% amplitude enhancement at least at one of the tested concentrations (Fig. 2G-S, Table 1). These compounds increased BMAL1:Luc2P amplitude in NIH3T3 cells without causing a significant deviation from the basal circadian period observed in DMSO-treated NIH3T3 cells at any tested concentration (SI Appendix, Fig. S2B).

**Figure 2:**
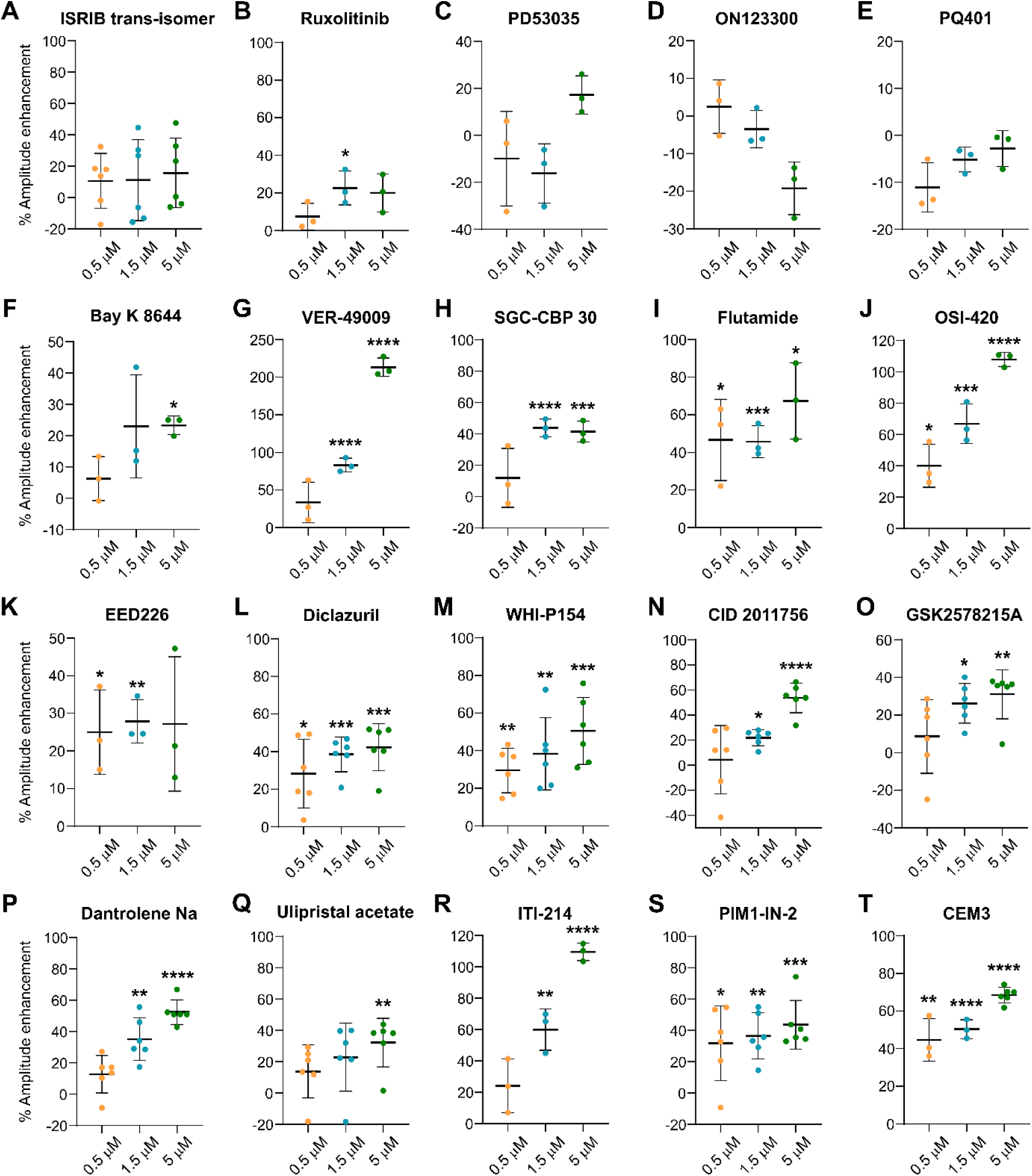
Validation of screening hits for Bmal1::Luc2P amplitude enhancement in NIH3T3 cells. (A-S) Quantification for change in Bmal1::Luc2P circadian rhythm amplitude above basal DMSO amplitude upon treatment of NIH3T3 cells with indicated concentrations of commercial stocks of 19 amplitude enhancer hits. Mean±SD, significance: between amplitude increase upon compound treatment vs DMSO; *P≤0.05, **P≤0.01, ***P≤0.001, ****P≤0.0001: statistical significance calculated with one-way ANOVA with two-stage Benjamini, Krieger, & Yekutieli correction. n=1-2 biological repeats, at least 3 technical replicates. (T) Same as (A-S) with positive control CEM3.

**Table 1:**
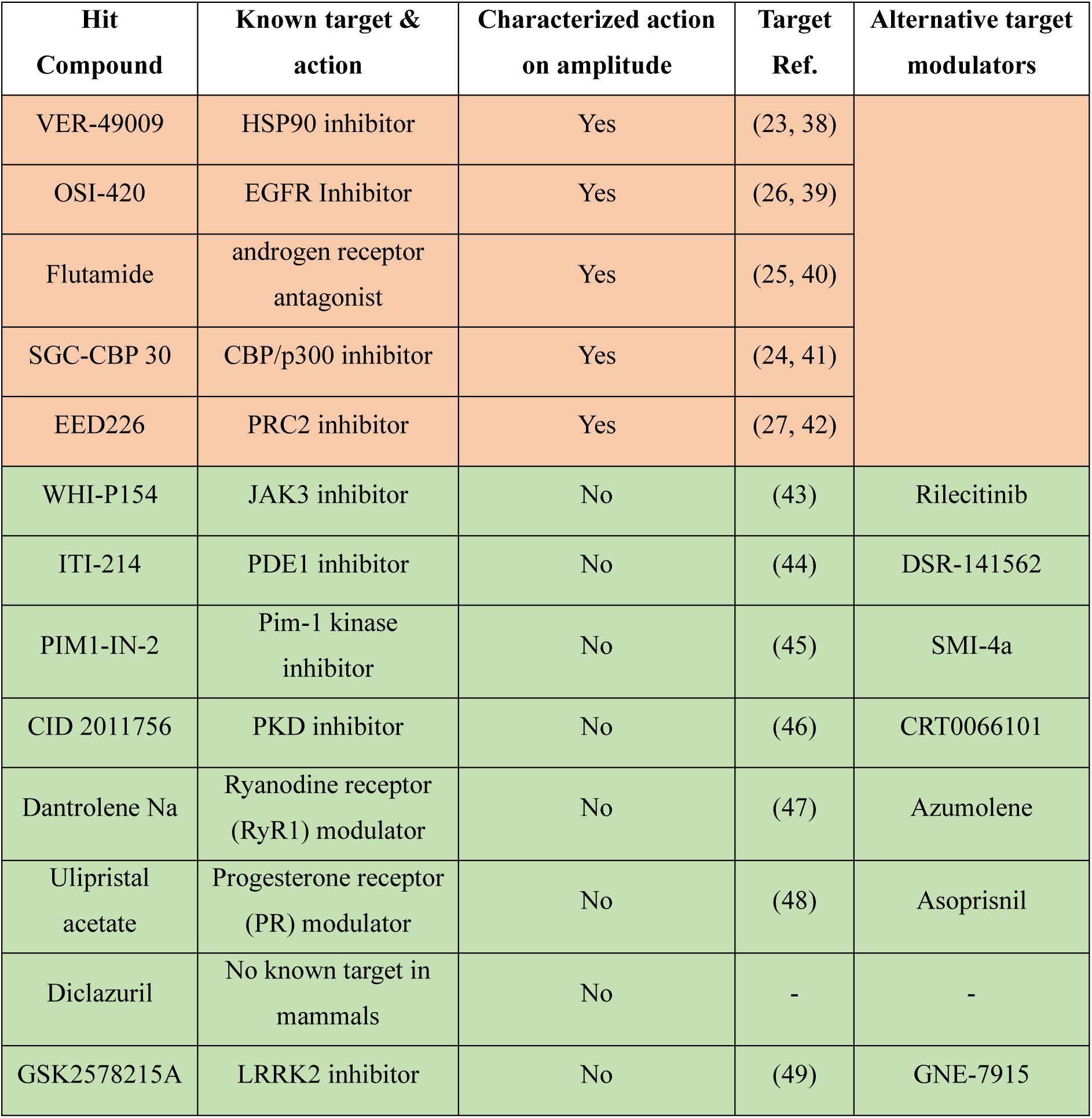
Validated screening hits, their targets, and alternative modulators.

Amongst these, protein targets of five compounds-VER-49009, SGC-CBP 30, Flutamide, OSI-420, and EED226 have been previously characterized as circadian amplitude modulatory proteins (Figure 2G-K, Table 1: in orange). Notably, eight compounds: WHI-P154, ITI-214, PIM1-IN-2, CID 2011756, Dantrolene Na, Ulipristal acetate, Diclazuril, and GSK2578215A, are novel amplitude enhancers as the compounds themselves and their targets have not been reported as modulators of molecular circadian rhythm (Fig. 2L-S, Table 1: in green). These compounds also did not significantly affect cell proliferation or death, as confirmed by the metabolic viability of compound-treated NIH3T3 cells (SI Appendix, Fig. S3).

Next, we validated the target-specific action of the amplitude-enhancing compounds by treating BMAL1::Luc2P-expressing cells with alternative inhibitors to the main published target of the novel compounds (Table 1, column 5). Except for CRT0066101 (an alternative inhibitor of PKD), all target modulators induced amplitude enhancement (Fig. 3). A dose-dependent amplitude response was observed with rilecitinib and azumolene (Figs. 3A and B, alternatives for WHI-P154 and Dantrolene Na, respectively), substantiating the role of their targets (JAK3 and RyR) in circadian amplitude modulation. Treatment with alternative inhibitors of LRRK2, PR, PIM1, and PDE1 (Figs. 3D-G) resulted in sustained concentration-independent amplitude enhancement.

**Figure 3:**
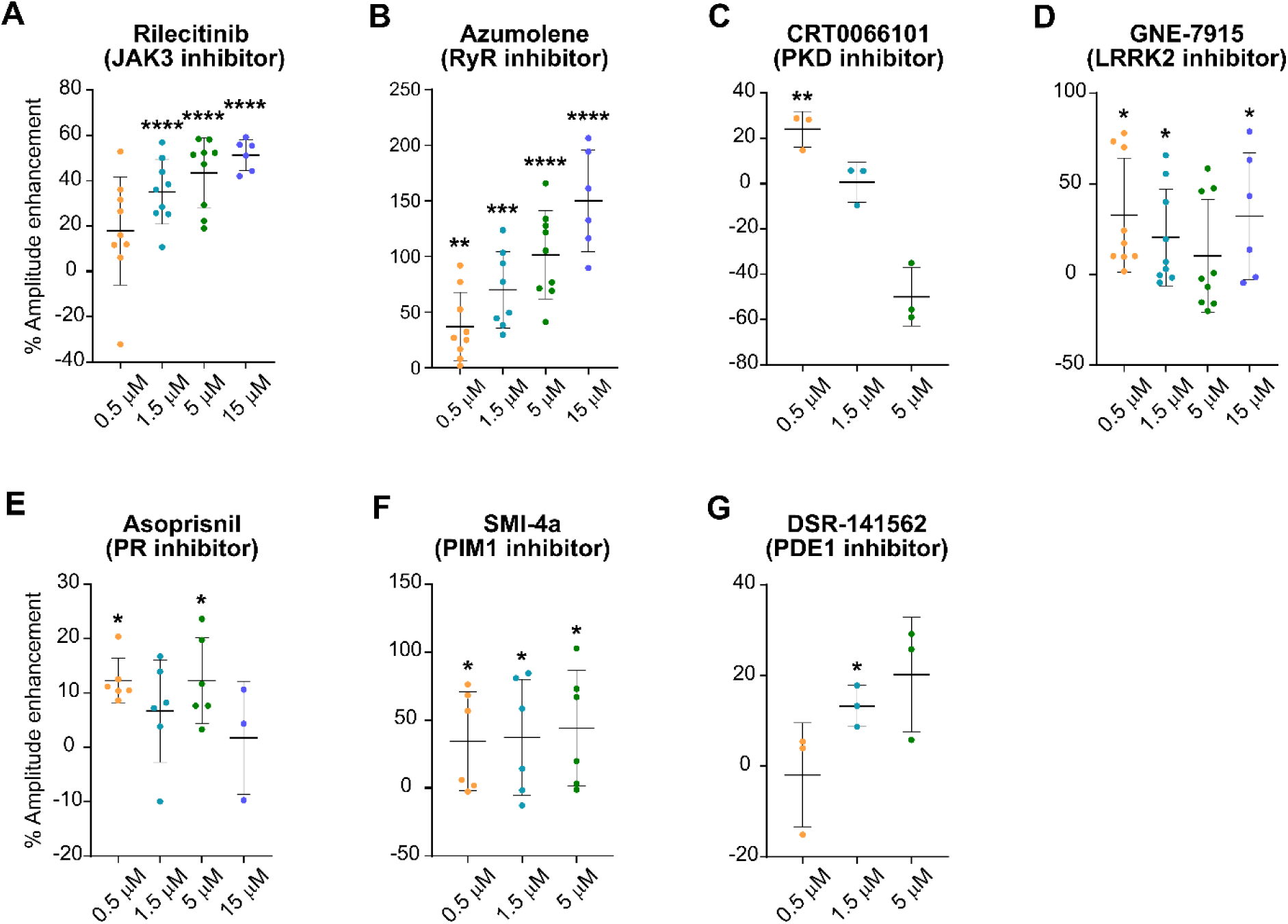
Target validation of screening hits with alternative inhibitors. (A-G) Quantification for change in Bmal1::Luc2P circadian rhythm amplitude above basal DMSO amplitude upon treatment of NIH3T3 cells with indicated concentrations of alternative inhibitors to targets of novel amplitude enhancer hits. Mean±SD, significance: between amplitude increase upon compound treatment vs DMSO; *P≤0.05, **P≤0.01, ***P≤0.001, ****P≤0.0001: statistical significance calculated with one-way ANOVA with two-stage Benjamini, Krieger, & Yekutieli correction. n=1-3 biological repeats, at least 3 technical replicates.

### Amplitude enhancement effects on a peripheral clock

After demonstrating the amplitude-enhancing effect of the compounds on the level of cell-monolayers, the next step towards studying their functional relevance was to examine if the hit compounds can also enhance molecular circadian rhythm on a higher level of organization, that is, animal-derived organotypic slices (20). Therefore, we tested whether the novel identified compounds exert an amplitude-enhancing effect on liver organotypic slices derived from PER2::LUC fusion protein expressing mice (21). To simultaneously validate the effects of multiple hits, mouse-derived acute slices were mounted on porous transwell inserts in a 24-well plate (Fig 4A), and bioluminescence was recorded for each slice using a plate reader over 3 cycles. The relatively higher organ volume of the liver and the amount of cells expressing the PER2::LUC construct enabled us to adapt the tissue-level circadian rhythm recording to mid-throughput drug-testing capabilities.

**Figure 4:**
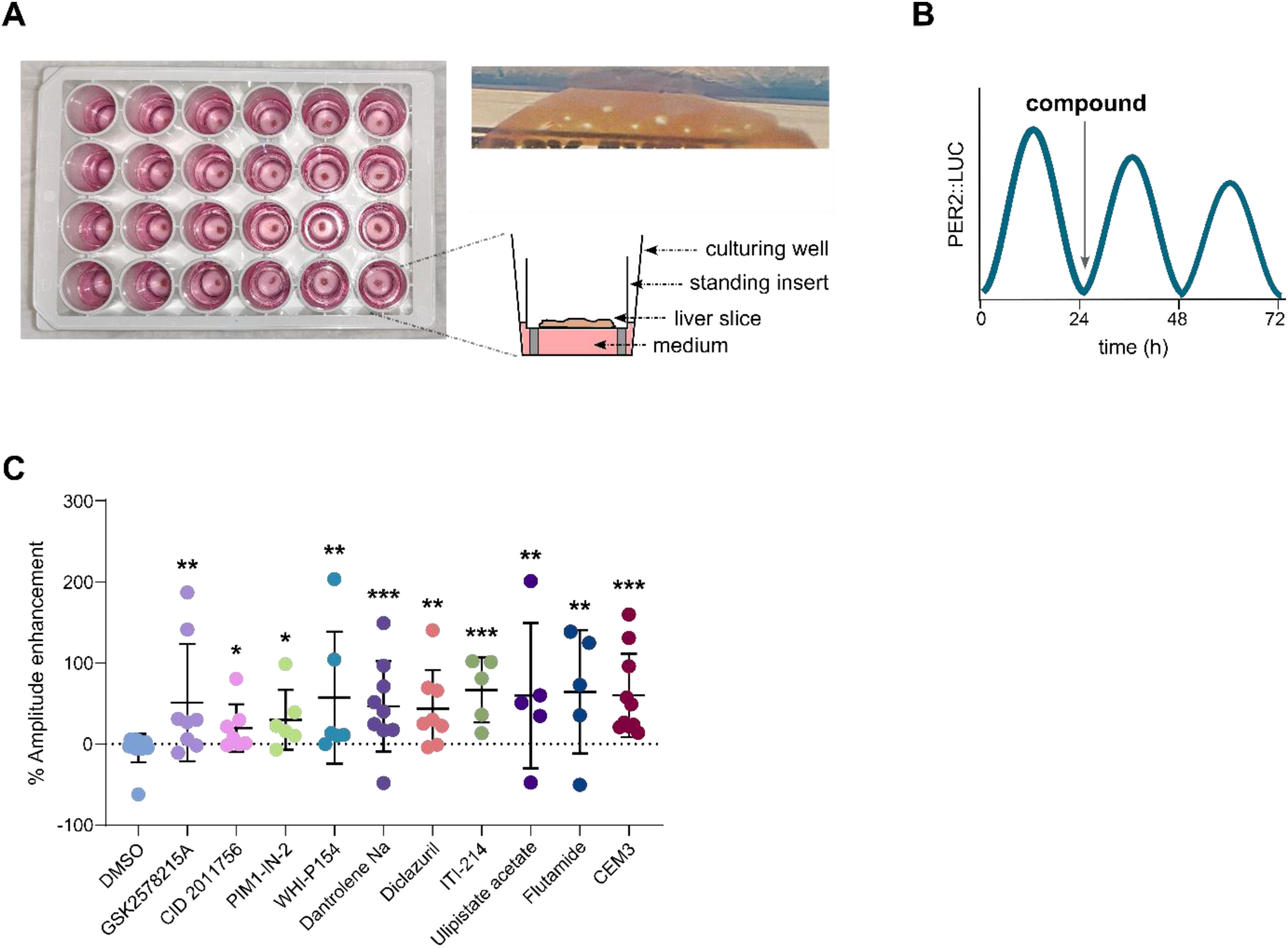
Effect of screening hits on PER2::LUC rhythm amplitude in liver-slice peripheral clock. (A) An image of a 24-well culture plate containing mouse-derived liver organotypic slices (left), a schematic of the side-view of a liver slice in the plate (bottom right), and an image of a liver cross-section sliced by Vibratome (top right). (B) Illustration of the molecular circadian rhythm to demonstrate the administration of hit compounds at the trough of the first (baseline) cycle. (C) Quantification for PER2::LUC circadian rhythm amplitude after application of the indicated hit compounds to liver slices at 30 µM. Mean±SD, each dot represents a single liver slice, *P≤0.05, **P≤0.01, ***P≤0.001: adjusted P-values for multiple comparisons, statistical significance calculated with Kruskal-Wallis test, with two-stage Benjamini, Krieger, & Yekutieli correction. n=4-10 liver slices.

As the acquired integrated bioluminescence signal can vary depending on the thickness, surface area, and viability of an individual slice, we applied compounds to slices after recording a baseline cycle and calculated the difference in amplitude between pre- and post-treatment cycles (Fig 4B). To compensate for the potential effect of the solvent, this difference in amplitude was normalized to the change induced upon DMSO treatment to derive relative amplitude enhancement. Known circadian amplitude enhancers of PER2 (CEM3 and Flutamide) showed significant amplitude enhancement on liver-slice tissue, thus validating our approach. Moreover, all hit compounds significantly enhanced the PER2::LUC amplitude of liver-slice clock (Figs. 4C, SI Appendix, Fig. S4), demonstrating their amplitude modulatory effect on the tissue level. With that, we also present a novel and physiologically relevant approach for screening modulators of circadian rhythms using animal-derived organotypic explants of liver in a bioluminescence plate reader.

### Amplitude enhancement effects on the central clock

Another crucial area that should benefit from amplitude-enhanced circadian rhythm is the SCN region of the brain. Notably, not all clock-modulating compounds reported in previous studies have shown effects in the SCN, despite their success in peripheral tissues. Therefore, we investigated whether the compounds identified in this study can enhance molecular circadian rhythm in the SCN-slices. We used SCN explants derived from PER2::LUC fusion protein expressing mice and measured the circadian rhythm in individual neurons by bioluminescence imaging of these brain organotypic slices (21). The top 6 hit compounds from the screen were applied to organotypic slices, and their effect on PER2::LUC rhythms was continuously recorded (Movie1, Fig. 5). To reduce variability between preparations, individual compounds were administered after recording a baseline cycle, and the difference in amplitude between pre- and post-application cycles was calculated to determine the net change in amplitude. Single-cell analysis of amplitude enhancement shows that the majority of SCN neurons showed increased amplitude of PER2::LUC rhythm after the application of compounds (Figs. 5A-C, SI Appendix, Fig. S5), in comparison to DMSO control. All the tested hit compounds significantly enhanced the PER2 circadian rhythm in SCN-slices, demonstrating their potential as central clock modulators (Fig. 5C).

**Figure 5:**
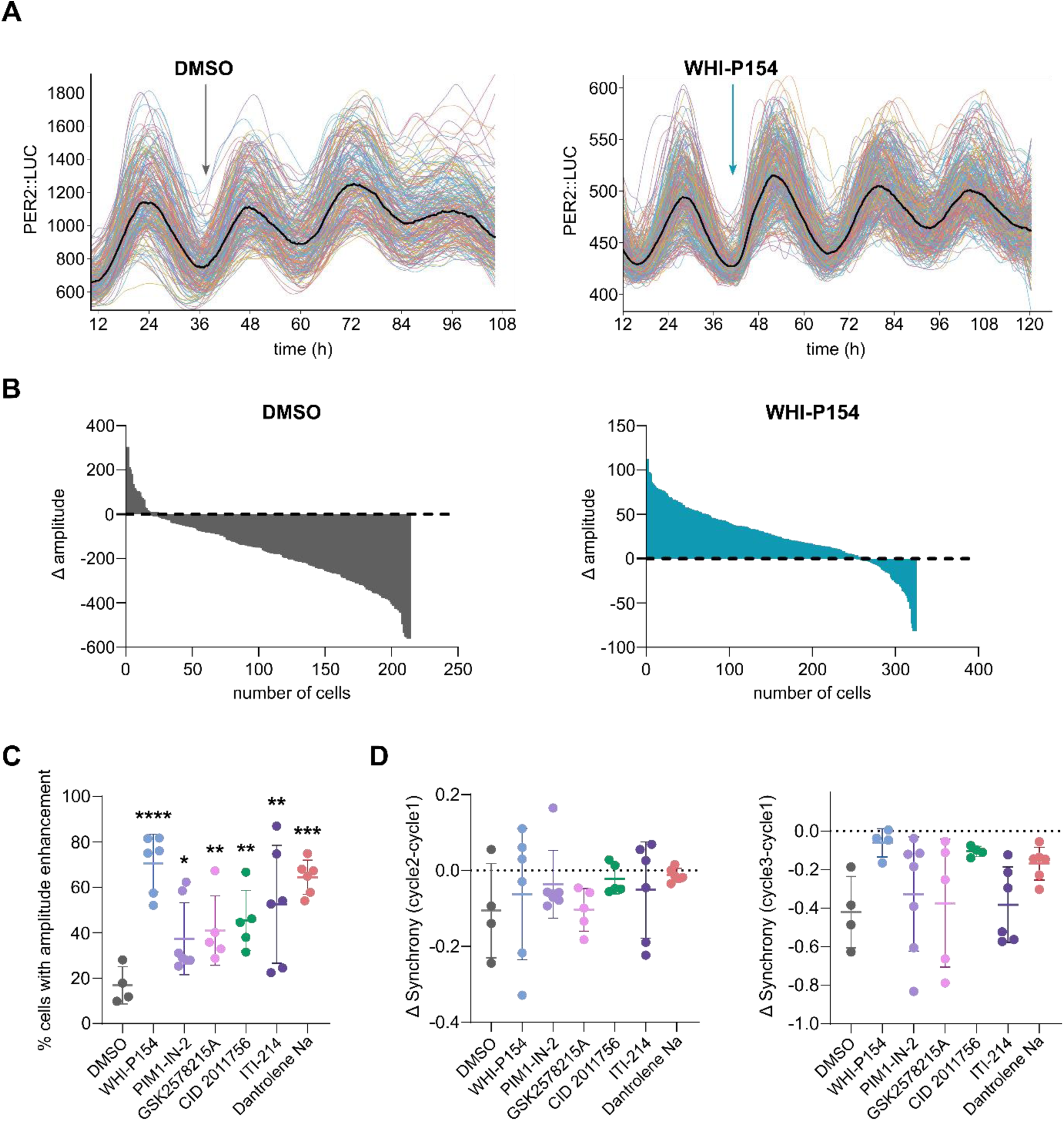
Effect of screening hits on PER2::LUC rhythm amplitude in neurons of the central clock. (A) Representative circadian rhythms of PER2::LUC expression in single cells from a suprachiasmatic nucleus (SCN) *ex vivo* explant obtained from a mouse brain, either after the application of vehicle (0.1% DMSO, left) or hit compound WHIP154 (30 µM, right). Individual light traces: single cell rhythms, solid black line: moving average of all traces from a slice. (B) Quantification of changes in the amplitude of PER2 rhythm in cells (from A). The difference in amplitude between before and after treatment with DMSO (left) or WHI-P154 (right) is plotted against cell number, sorted by the magnitude of the induced change in amplitude upon compound treatment. The point of intersection of the graph with the dashed line at Δ amplitude = 0 marks the transition from cells showing enhanced amplitude to cells with decreased amplitude of their rhythms after treatment. (C) Average fraction of cells per animal that show an increased amplitude after application of the indicated hit compounds to SCN-slices at 30 µM. Mean±SD, *P≤0.05, **P≤0.01, ***P≤0.001, ****P≤0.0001: adjusted P-values for multiple comparisons, statistical significance calculated with one-way ANOVA with two-stage Benjamini, Krieger, & Yekutieli correction. n=4-7 animals. (D) Difference in synchrony per SCN explant before and after treatment with indicated hit compounds to SCN-slices at 30 µM. Left: change in synchrony between the second and the first cycle, right: change in synchrony between the third and the first cycle. Mean±SD, no significant comparisons: statistics calculated with one-way ANOVA with two-stage Benjamini, Krieger, & Yekutieli correction, n=4-7 animals.

Here, the amplitude enhancement on the tissue level can either be due to amplification of clock protein rhythms on the single-cell level or by increasing phase synchrony between them (22). To understand the mechanism behind amplitude enhancement, we calculated the induced difference between synchronization before (cycle 1) and after DMSO or compound treatment (cycle 2 and cycle 3). The degree of synchronization did not significantly change upon any of the compound treatments in comparison to DMSO (Fig 5D). Therefore, amplitude enhancement induced by WHIP154, PIM1-IN-2, GSK2578215A, CID2011756, ITI-214, and Dantrolene Sodium was achieved through the enhancement of clock protein rhythm in individual cells, instead of increasing synchrony between cells.

## Discussion

In this study, we report the identification of eight compounds: WHI-P154, ITI-214, PIM1-IN-2, CID 2011756, Dantrolene-Na, Ulipristal acetate, Diclazuril, and GSK2578215A as novel circadian rhythm amplitude enhancers by screening a drug-repurposing library of 5,631 compounds. These compounds were initially characterized by performing a high-throughput screen in Bmal1-luciferase fibroblasts, followed by validation in liver and SCN tissues from PER2-luciferase expressing mice. We focused on circadian amplitude, a comparatively underexplored yet physiologically important parameter. Previous compound screens have identified a limited number of circadian amplitude enhancers, with only a few demonstrating sustained effects across multiple oscillatory cycles and reporter systems without significant alterations to circadian period. Moreover, amplitude enhancers with activity across both central and peripheral clocks remain uncommon. Notably, in this study, we find eight robust amplitude-enhancing molecules with consistent effects across both peripheral and central clock tissue, highlighting their potential as broadly applicable modulators of circadian function. As damped circadian amplitude has been an early-stage hallmark of metabolic disorders, neurodegenerative disorders, and ageing, identification of this array of amplitude-enhancing compounds can lead to the development of novel chronotherapeutic approaches to counter the progression of these conditions (6).

Amongst the published amplitude-enhancing molecules, CEMs (e.g., CEM3) have shown excellent potential to enhance amplitude across tissues. However, their intracellular targets and mechanism of action are still unresolved (17, 22). For Nobiletin, although the intracellular targets and applicability for metabolic disorders have been proposed, the amplitude enhancement is restricted to specific peripheral tissues (11). Thus, there remains a need to identify new circadian-modulating molecules and their respective targets. Such discoveries could enable the rational development of more potent and selective inhibitors, while also facilitating tissue-specific therapeutic applications in organs where the target plays an important physiological role. Another important consideration is that many reported amplitude enhancers do not maintain their effects across multiple oscillatory cycles. To identify compounds capable of producing sustained amplitude enhancement, we applied wavelet-based analysis for cell rhythms. This approach revealed that all validated compounds produced a sustained increase in time-resolved circadian amplitude across multiple oscillatory cycles.

The targets of compounds in this study (Table 1) can also be valuable for understanding previously unidentified signaling pathways to modulate circadian amplitude. However, a known main target of a compound may not be the one responsible for the observed effect on the circadian rhythm, or the activity of a compound can be through more than one target. We verified this by testing the effect of alternative inhibitors of the known targets of our hits. We found dose-dependent enhancement of circadian rhythm amplitude upon inhibition of JAK3, Ryanodine receptor, and phosphodiesterase I, confirming the involvement of these targets in rhythm modulation. However, the involvement of the published targets of the remaining hits might still be ambiguous. Inhibition of LRRK2, progesterone receptor, and PIM1 showed a sustained dose-independent amplitude enhancement. This can either mean that the maximum response is already reached at the lowest tested dose, or that there might be additional targets of the hit compounds that are involved in inducing the observed amplitude enhancement response associated with the compound. Additionally, one of the hits, Diclazuril, an antiprotozoal agent, does not have known mammalian targets, making it a promising entry point for identifying novel circadian regulators.

Five compounds and/or their targets identified in this screen have been previously reported as circadian amplitude modulators, providing independent validation for the strength of our screening methodology. This includes HSP90 (target of VER-49009), CBP/p300 (target of SGC-CBP30), PRC2 (target of EED226), androgen receptor (target of flutamide), and EGFR (target of OSI-420), which have been shown to modulate circadian amplitude in specific cell types and tissues (23–27). For the eight newly identified amplitude modulators, the molecular mechanisms through which target engagement leads to enhanced circadian amplitude remain largely unresolved. Target modulation may influence several regulatory mechanisms, including protein-protein interactions and post-translational modifications of core clock proteins, which could enhance the positive arm of the TTFL and contribute to increased circadian amplitude.

For example, inhibition of PDE1 has been shown to increase cAMP levels, thereby enhancing CREB phosphorylation and promoting *Per2* transcription (28). Progesterone receptor (PR), a nuclear receptor, has tissue-dependent mechanisms that can link it to the clock by recruiting CBP/p300, which can repress the formation of CLOCK/BMAL1 transcription activator heterodimer (24, 29). It can also cross-talk with ERK, cAMP, and Ca^2+^ signaling to regulate clock protein expression (30). Serine/threonine kinases, including PKD and PIM1, may regulate circadian protein stability, expression, and interactions through phosphorylation of multiple clock components, including BMAL1, CLOCK, PER, and CRY; however, direct evidence linking these kinases to circadian amplitude regulation remains limited. For the ryanodine receptor, there is ambiguity about how it might regulate the amplitude of BMAL1 and other clock proteins, despite the strong evidence of BMAL1 regulating the circadian rhythm of the ryanodine receptor. One possibility is through the regulation of Ca^2+^ signaling and CREB-mediated transcription, both of which can regulate clock gene transcription (31). For JAK3 and LRRK2, the molecular mechanisms connecting their activity to circadian clock regulation remain largely unexplored. However, their established roles in disease-associated pathways suggest potential links to circadian dysfunction. JAK3 is involved in inflammatory signaling, and proinflammatory transcription factors such as NF-κB have been shown to suppress CLOCK/BMAL1 transcriptional activity (32). Therefore, JAK3 inhibition may indirectly influence clock gene expression through modulation of inflammatory pathways. Similarly, reduced BMAL1 expression has been causally associated with oxidative stress and synaptic dysfunction in Parkinson’s disease, a disorder in which LRRK2 plays a major role (33). However, whether and how LRRK2 and BMAL1 regulate each other in the context of Parkinson’s disease remains unknown. Together, these findings highlight the potential disease relevance of circadian amplitude modulation and underscore the need for further investigation into the mechanisms linking these targets to the molecular clock.

Tissue-specific expression patterns and signaling context can influence how small molecules modulate circadian rhythms in central and peripheral clocks. To evaluate this, we validated their activity in organotypic liver and SCN slice cultures. We used a mouse model expressing the PER2::LUC fusion protein, which provides a real-time readout of the negative limb of the TTFL and enables assessment of whether transcriptional changes observed at the cell level are reflected in the protein oscillation level. Notably, all compounds enhanced circadian amplitude in both tissues, demonstrating their conserved effects across distinct physiological contexts. Single-cell imaging of SCN slices further revealed that amplitude enhancement arises from increased oscillatory strength within individual cells rather than changes in intercellular synchronization. This suggests that the identified compounds primarily act through cell-autonomous mechanisms to reinforce molecular clock function. Together, the identified compounds highlight the robustness of circadian amplitude modulation across tissues and provide a foundation for pharmacological strategies to target both central and peripheral clocks by drug repurposing.

## Materials and methods

### NIH3T3 cell culture

NIH3T3 cells were obtained from ATCC (American Type Culture Collection). Cells were grown and maintained in Dulbecco’s Modified Eagle’s Medium (DMEM; 11995065, Gibco) supplemented with 10% (v/v) Fetal Bovine Serum (FBS), 1% L-glutamine, 1% penicillin-streptomycin at 37°C and 5% CO_2_. Cells were subcultured twice a week on 10 cm Ø plates.

### Live-cell circadian rhythm assay with NIH3T3 fibroblasts

NIH3T3 cells were seeded at 1.5×10^5^ cells/well in a 6-well plate (3 cm Ø) and cultured until they reached 70% confluency. Cells were transfected with 2 µg per well of BMAL1 promoter-inserted pGL4.11 plasmid (9PIE666, Promega) using X-tremeGENE™ HP (Merck) as per the manufacturer’s protocol. The cells were incubated for 18 hours at 37°C, 5% CO_2,_ before being transferred to white-walled, clear-bottom 384-well (6007480, ViewPlate-384 TC, Revvity, MA, USA) or 96-well assay plates (6005181, ViewPlate-96, Revvity) at a seeding density of 1.2×10^4^ or 4×10^4^ cells/well, respectively, followed by further incubation for 24 hours. Two hours before administration of the compounds, cells were synchronized with 5 µM Forskolin (TRC-F701800, LGC standards, Germany).

Compound dispensing and rhythm recording for high-throughput screening experiments: Culturing medium was removed from the 384-well assay plate containing cells and replaced with 20 µL CO_2_-independent medium (18045088, ThermoFisher, MA, USA) supplemented with 10% FBS, 1% penicillin-streptomycin and 0.2 mM D-Luciferin (15285723, Thermo Scientific) with a Biotek MultiflowFX liquid handler. Next, compounds were dispensed from library plates (384-well) to the 384-well assay plate using a Labcyte Echo650 acoustic dispenser (Beckman Coulter, CA, USA), followed by redispensing of 30 µL CO_2_-independent medium. Known circadian modulators CEM3 and JNJ-6204 were used as positive controls for amplitude enhancement and period lengthening, respectively, and DMSO was used as a vehicle control. For the primary 4-timepoint screening of 5,631 compounds, the percentage of DMSO was maintained at 0.1%, and for the subsequent screenings with continuous recording, it was at 0.5% across all the treated wells.

The 384-well plate containing compound-treated cells was sealed with adhesive aluminum foil and loaded into a PHERAstar FSX plate reader (BMG Labtech, Germany) maintained at 37°C to record bioluminescence over 3.5 days at a measurement interval of 10 mins. Bioluminescence was measured with the following plate reader settings: LUM optic module, bottom optics, focal height 4.3 mm, 96/384 aperture spoon for crosstalk reduction, and orbital averaging at 2 mm well diameter.

Compound dispensing and rhythm recording for low-throughput experiments: Compounds were dispensed using a Labcyte Echo650 acoustic dispenser to a blank 96-well destination plate (651201, Greiner) from an Echo-ready 384-well source plate (Beckman Coulter) containing 10 mM stocks of compounds in DMSO. Compounds were then diluted to the assay concentrations in 200 µL CO_2_-independent medium with the composition mentioned above and transferred manually to the 96-well assay plate containing cells. The percentage of DMSO was maintained at 0.1% across all the treated wells. The plate was sealed with adhesive aluminum foil and loaded into a PHERAstar FSX plate reader (BMG Labtech, Germany) maintained at 37°C to record bioluminescence over 3.5 days at a measurement interval of 10 mins. Bioluminescence was measured with the following plate reader settings: LUM optic module, bottom optics, focal height 4.4 mm, 96/384 aperture spoon for crosstalk reduction, and orbital averaging at 5 mm well diameter.

### Analysis of BMAL1::Luc2P circadian rhythm data

For primary screening where BMAL1:::Luc2P bioluminescence was recorded at 4 time points (6 hrs, 10 hrs, 24 hrs, and 30 hrs), the amplitude increase induced at each sampled time point (t) by compound stimulus was calculated by normalizing it to the average (*DMSO_Avg_*) and standard deviation (*DMSO_SD_*) in DMSO-treated samples:

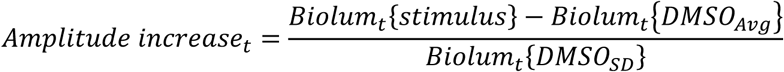

A compound was defined as a hit if the above calculated amplitude increase at each sampled time point is above 1 at timepoints 6 hrs, 24 hrs and 30 hrs, as well as above 1.5 at 10 hrs.

For subsequent screenings where a ∼3.5-day-long circadian bioluminescence rhythm was recorded, the time-series data were analyzed using pyBOAT (Python-based Oscillatory Analysis Tool) to estimate the rhythm parameters: phase, period, and amplitude (34). The data were detrended using Sinc filter with a cut-off period of 48 hrs. Signals were then subjected to continuous Morlet wavelet transformation, with the following settings: smallest period 2 hrs, number of periods 1000, highest period 36 hrs. The dominant circadian component was identified by extracting the wavelet ridge, defined as the period of maximal wavelet power at each time point. For each series, ridge-associated parameters, including instantaneous period, amplitude, and wavelet power, were exported for downstream analysis. The power threshold was set to a constant value of 20 across the whole dataset (∼15-20% of the average maximum ridge power observed) to exclude low-confidence oscillatory estimates and exclusively retain the time points within the high-power rhythmic phase.

Exported ridge data were further processed using custom Python scripts. Instantaneous phase was reconstructed by integrating the ridge-derived period over time to define circadian cycles. Individual circadian cycles were distinguished by successive increases of 2π in cumulative phase. For each complete cycle, mean and peak ridge amplitude, mean period, and average wavelet power were calculated. In addition, mean ridge amplitude, period, and power were computed across all included timepoints for each series. Based on these estimated parameters, individual criteria were defined for period and amplitude to find hits in each round of screening (ref. supplementary data and scripts)

### Cell viability assay

Metabolic viability of NIH3T3 cells was assessed using the CellTiter-Blue® Cell Viability Assay (G8081, Promega) according to the manufacturer’s instructions. Cells were seeded in COSTAR 96-well plates (3599, Corning) at a density of 2000 cells per well in 200 µL complete growth medium and allowed to adhere overnight at 37°C in a humidified incubator with 5% CO₂. The following day (upon 50% confluency), cells were treated with amplitude enhancer hits at 5 µM. After 24 hours, CellTiter-Blue® reagent was added to each well at 20% of the culture volume (e.g., 20 µL reagent per 100 µL medium) and plates were incubated for 1 hr at 37°C. Fluorescence was measured using a CLARIOstar plate reader (BMG Labtech, Germany) with excitation at 560 nm and emission at 590 nm. Cell viability was calculated as a percentage relative to DMSO-treated control wells, which were set to 100%.

### Animals

Male homozygous PER2::LUC knock-in mice were used in this study. All cells in these animals express a modified clock protein PERIOD 2 (PER2) fused to luciferase (PER2::LUC) (21). This enables the measurement of the circadian rhythms of the PER2 protein in single cells using bioluminescence imaging. Mice (3-6 months old) were bred and housed at the Leiden University Medical Center animal facility in climate and light-controlled cabinets (23°C; 12h light, 12h dark) with access to food and water *ad libitum.* The experiments were performed in accordance with the Dutch law on animal welfare and have been approved by the animal experiments committee Leiden (PE.20.041.010).

### Bioluminescence circadian rhythm assay with organotypic liver slices

Animals were sacrificed 5 h before lights-off (Zeitgeber Time (ZT) 7 with ZT0 defined by lights-on), livers were extracted, and slices were prepared as described previously (20, 35). The left lateral liver lobe was excised and trimmed to obtain an 8×16 mm cube. This cube was incubated for 30 minutes on ice in carbogen (95% O_2_-5% CO_2_)-bubbled Krebs-Henseleit Buffer (KHB) (K3753, Sigma-Aldrich) reconstituted with 2.5 mM CaCl_2_ and 23.8 mM NaHCO_3_. Next, the cube was embedded in 4%, low-temperature-gelling agarose (Sigma-Aldrich A4018) and left to harden on ice for 5 minutes. The surrounding agar was trimmed and glued with cyanoacrylate (Loctite 401, Henkel) to the vibratome chamber. The chamber was placed in the Vibratome (VT1000S, Leica) and filled with cold KHB, continuously bubbled with carbogen. Individual liverslices of 250 µM thickness were transferred onto a nylon mesh of a slice holder situated in a beaker containing carbogen-bubbled KHB just covering the slices, and incubated for 20 minutes at 4° C to reduce bile. A total of 24 organotypic slices of similar size (2×2 mm) were obtained from an individual liver cube. The slices were transferred to a 24-well imaging plate (1450-603, VisiPlate-24 TC, Revvity) with permeable membrane insert (PICM01250, Millicell, Merck) and cultured in 350 µL medium at 305 mOsm consisting of DMEM (SLM-241, Merck), 1% penstrep, 1% amphotericin, 1% non-essential amino acids, 5 µg/mL insulin, 10 mM HEPES, 50 μg/mL ascorbic acid, with added 0.2 mM D-Luciferin (15285723, Thermo Scientific).

The transparent bottom plate containing 24 liver slices was sealed with adhesive aluminum foil and loaded into a PHERAstar FSX plate reader (BMG Labtech, Germany) maintained at 37°C to record PER2::LUC expression over 3 days at a measurement interval of 10 mins. Bioluminescence was measured with the following plate reader settings: LUM optic module, bottom optics, focal height 3 mm, and spiral scanning at 14 mm well diameter. The initial cycle was recorded as a baseline cycle for each slice; the compounds were administered at the trough of this cycle, and the recording was continued. The obtained bioluminescence oscillations for each stimulus condition were smoothed using a rolling average over a 2-hour window, with a second-order polynomial regression in GraphPad Prism 8.4.2. Peaks and troughs were detected in each circadian cycle for each stimulus, and the rhythm amplitude was calculated by subtracting the trough bioluminescence value from the peak (Fig. 1A). The percentage increase in the amplitude induced by a compound stimulus was estimated by normalizing the post-stimulus (cycle 2) amplitude to the pre-stimulus basal amplitude (cycle 1) using the following formula:

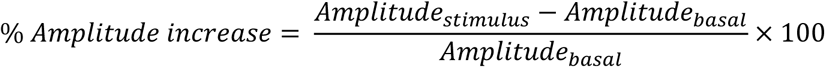

To correct for the effect of solvent DMSO on the rhythm, the average amplitude change induced by DMSO was subtracted from the amplitude increase for each slice, for each experiment, using the following formula:

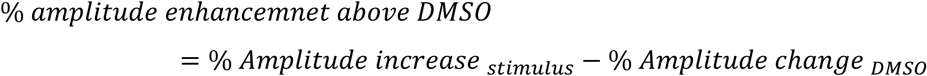

### Bioluminescence imaging of organotypic SCN explants

Organotypic explants of the SCN were prepared as described previously (22, 36). Briefly, animals were sacrificed 5 h before lights-off (Zeitgeber Time (ZT) 7 with ZT0 defined by lights-on). Hypothalamic slices (200 μm thick) containing the SCN were prepared using a vibrotome (VT 1000S, Leica) from rapidly excised brains in ice-cold modified artificial cerebrospinal fluid containing 116.4 mM NaCl, 5.4 mM KCl, 1.0 mM NaH_2_PO_4_, 0.8 mM MgSO_4_, 1.0 mM CaCl_2_, 4.0 mM MgCl_2_, 23.8 mM NaHCO_3_, 15.1 mM D-glucose, and 5 mg/L gentamicin (Sigma-Aldrich, Munich, Germany) saturated with 95% O_2_-5% CO_2_ and buffered to pH 7.4. SCN explants (ca 1×1mm) were cut from one selected slice per animal and placed on a Millicell membrane insert (PICM0RG50, Merck-Milipore, Burlington, MA) in a custom-made perfusion/imaging chamber filled with 1.7 mL Dulbecco’s Modified Eagle’s Medium (D7777, Sigma-Aldrich) supplemented with 10 mM HEPES buffer (Sigma-Aldrich), 2% B-27 (Gibco, Landsmeer, the Netherlands), 5 U/mL penicillin, 5 μg/mL streptomycin (0.1% penicillin-streptomycin; Sigma-Aldrich), and 0.2 mM D-luciferine sodium salt (Thermo Scientific Pierce Cat.#88293). Push-pull syringes were connected via tubing, and the application syringe was pre-filled with medium containing either the hit compound (30 μM) dissolved in DMSO (0.1%) or only DMSO (0.1%).

The perfusion chamber was mounted on a fixed-stage upright microscope (BX51WIF, Olympus) in a light-tight, temperature-controlled enclosure (Life Imaging Services, Reinach, Switzerland) kept at 37°C. Microscope position and focus were controlled by a motorized stage (XY-shifting table 240, Luigs & Neumann, Ratingen, Germany) and motorized focus (MA-42Z, Märzhäuser, Wetzlar, Germany). Bioluminescent images from two SCN explants from different animals were acquired by a cooled CCD camera (Retiga LUMO, Teledyne Photometrics, Birmingham, United Kingdom) using an exposure time of 29 minutes, with a 1-h sampling interval for 6 days. Application of either DMSO or compounds was performed after the first complete cycle of PER2 rhythms, 8 hours before the second peak (ZT4). Image acquisition was controlled by Micro-Manager ImageJ software (University of California, San Francisco) (37).

### Image analysis of organotypic SCN explants

A custom-made MATLAB-based (Mathworks, Natick, MA, USA) algorithm was used to analyze the time series of bioluminescence images, as described previously (36). Briefly, cell-like regions of interest (ROI) were defined using peak-intensity maps created from the images. Average intensity values for each ROI were calculated, which resulted in traces of bioluminescence intensity representing the PER2::LUC expression rhythms of single SCN cells. For analysis of peak time, raw PER2::LUC expression traces were smoothed and resampled to one data point per minute. Amplitude was calculated as the difference between the trough and the peak of the PER2::LUC rhythm. Changes in amplitude due to compound application were calculated between the first (untreated) cycle and the first cycle after treatment. All values for amplitude changes were normalized to the average change in amplitude after application of the control medium containing DMSO. The synchrony of PER2::LUC rhythm phases of single SCN cells was calculated using the Kuramoto order parameter (r) with possible values between 0 (fully unsynchronized) and 1 (fully synchronized).

## Data Availability

All data is available in the figures and source data files. Custom Python scripts used for analyzing and plotting the screening data can be found at https://github.com/MaiJoshi4/circadian_amplitude_HTS

## Supporting information

MovieS1_SCN

## Acknowledgements and funding

We thank Ines Chaves (Erasmus MC, Rotterdam, The Netherlands) for providing the Bmal1:Luc2P plasmid, Inge Snijders and Laura Heitman (LACDR, Leiden University, The Netherlands) for providing CEM3, and the Oncode Institute for providing the drug repurposing library. This work was supported by the BioClock consortium funded by the research program NWA-ORC by the Dutch Research Council (NWO), and by Oncode Accelerator, a Dutch National Growth Fund project under grant number NGFOP2201.

## Author Contributions

M.S.J., S.H.M., and P.P.G. designed research; M.S.J and F.P.A. performed experiments; M.S.J. analyzed data; P.P.G., S.H.M and J.H.M. provided supervision; M.S.J, J.H.M, S.H.M and P.P.G wrote the manuscript.

