## Supplemental figures and tables for "Identification of small-molecule enhancers of circadian rhythm amplitude in central and peripheral clocks"

* Correspondence: **Maitreyi S. Joshi**

**This PDF file includes:**

Figures S1 to S5
Table S1
SI: Molecular structures of identified hit compounds
Legends for Movies S1
Legends for Datasets S1 to S5

**Other supporting materials for this manuscript include the following:**

Movie S1
Datasets S1 to S5

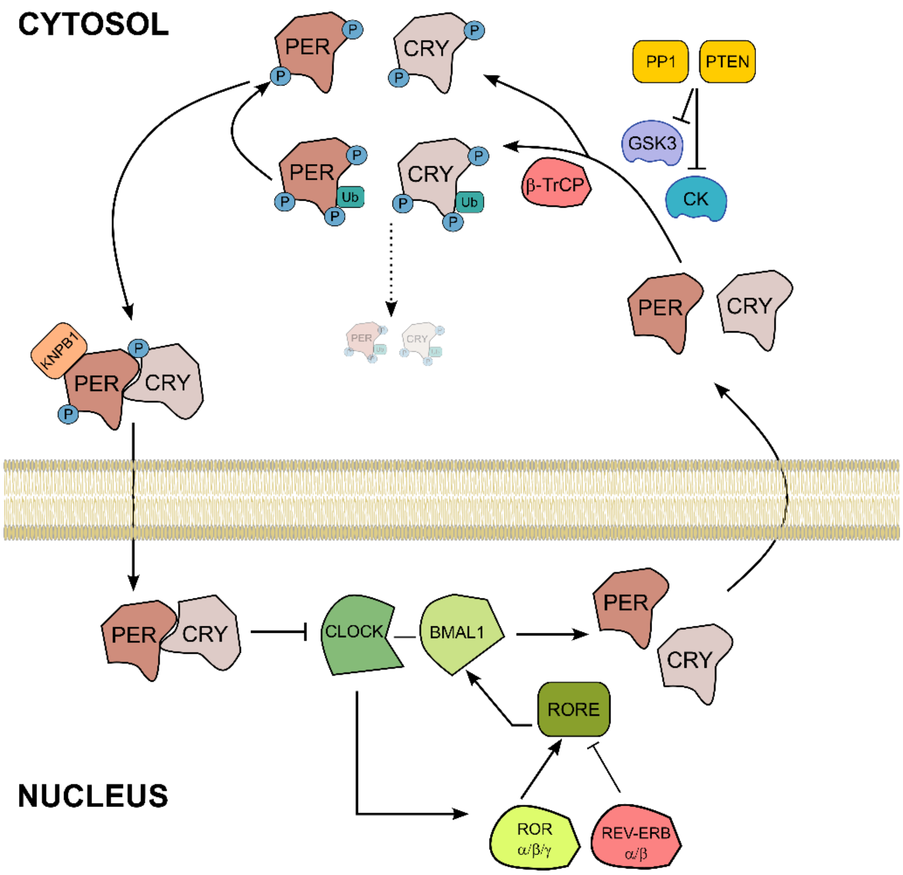


**Fig. S1: Molecular architecture of circadian rhythm network**

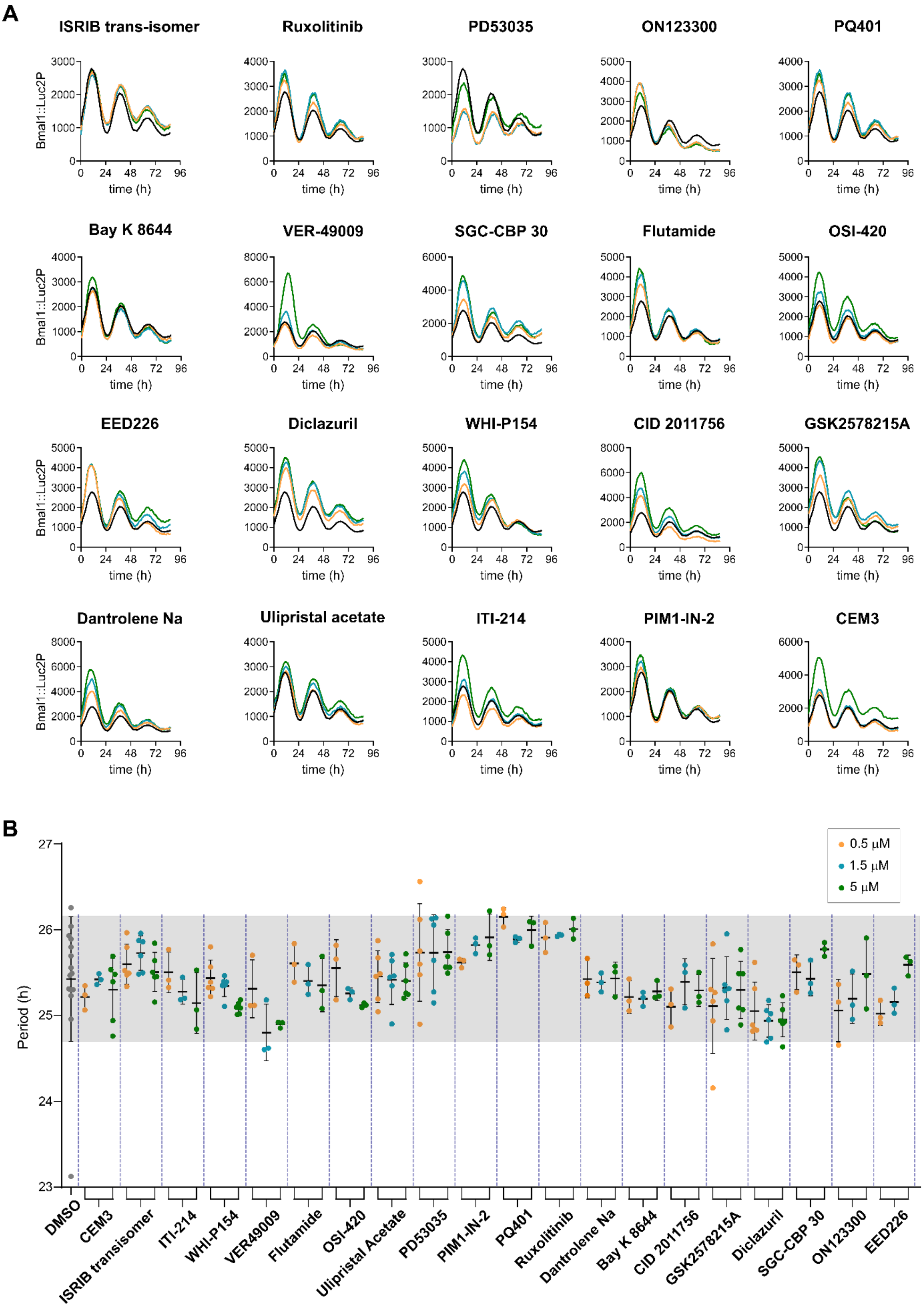


**Fig. S2: Bmal1::Luc2P circadian rhythm and period distribution for screening hits**

(A) Representative Bmal1::Luc2P bioluminescence profiles from NIH3T3 cells upon treatment with screening hits at indicated concentrations.

(B) Circadian rhythm period of Bmal1::Luc2P oscillations upon treatment with screening hits at indicated concentrations. The shaded grey area represents the standard deviation in circadian period upon vehicle (DMSO) treatment. Mean±SD, n=1-2 biological repeats and at least 3 technical replicates.

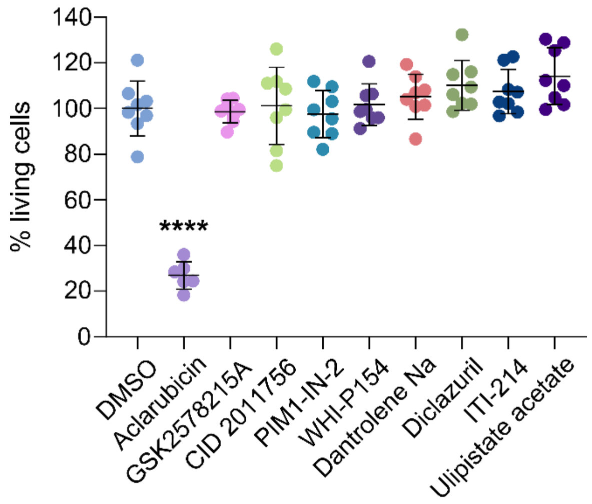


**Fig. S3: Effect of screening hits on cell viability**

The percentage of living NIH3T3 cells after 24 h of treatment with the indicated screening hits at 5 µM. The fraction of metabolically viable cells is estimated from fluorescence readout (resazurin-to-resorufin conversion) and normalized to the effect induced by DMSO. Mean±SD, ****P≤0.0001: adjusted P-values for multiple comparisons, statistical significance calculated with one-way ANOVA with Dunnett correction. n=3 biological repeats and 6-8 technical replicates.

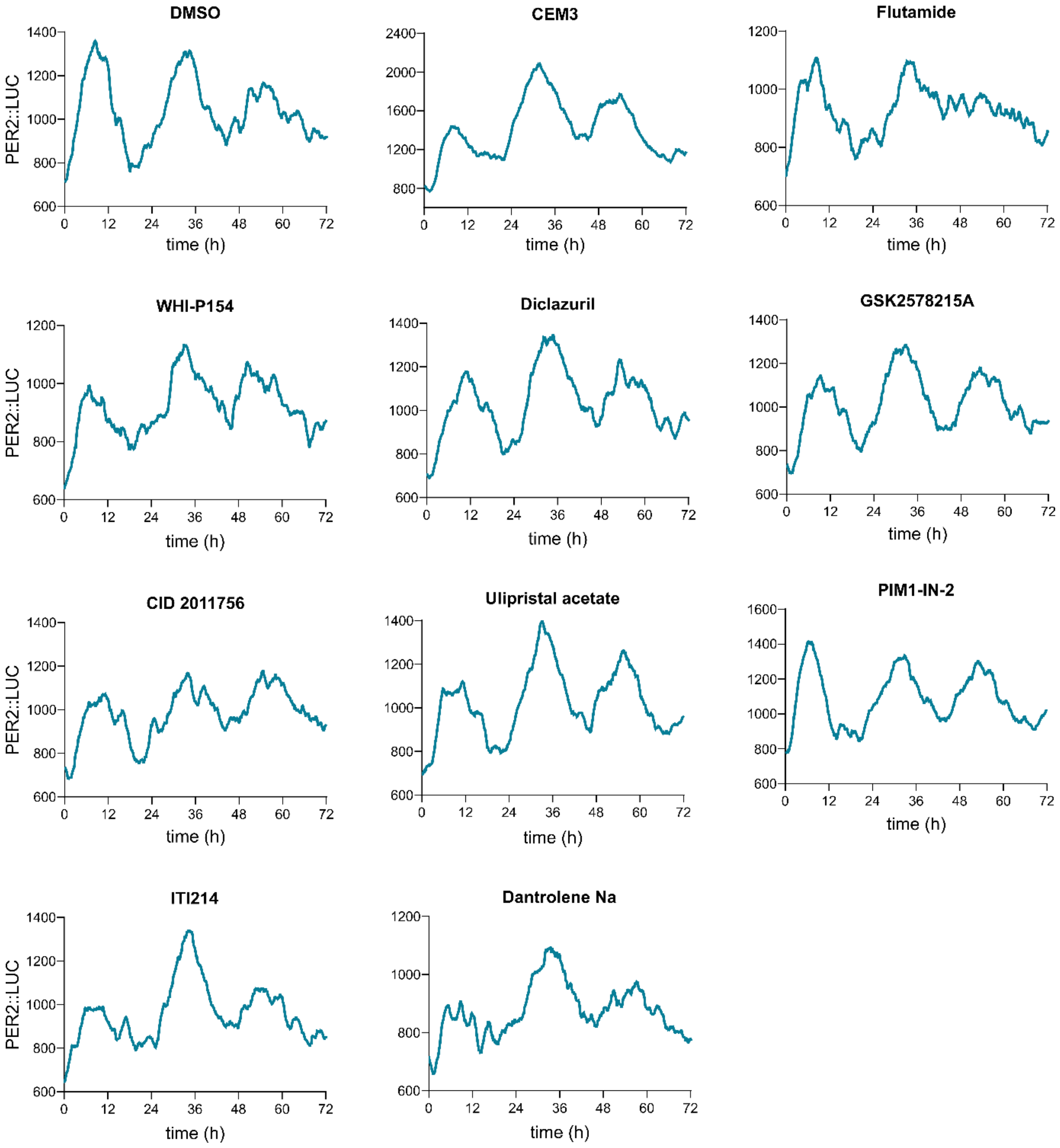


**Fig. S4: Effect of hits on PER2::LUC rhythm amplitude in liver-organotypic slices**

Representative circadian rhythms of PER2::LUC expression recorded as bioluminescence from acute slices of mouse liver. Slices were treated with vehicle (0.1% DMSO) or hit compounds (30 µM) after the first cycle.

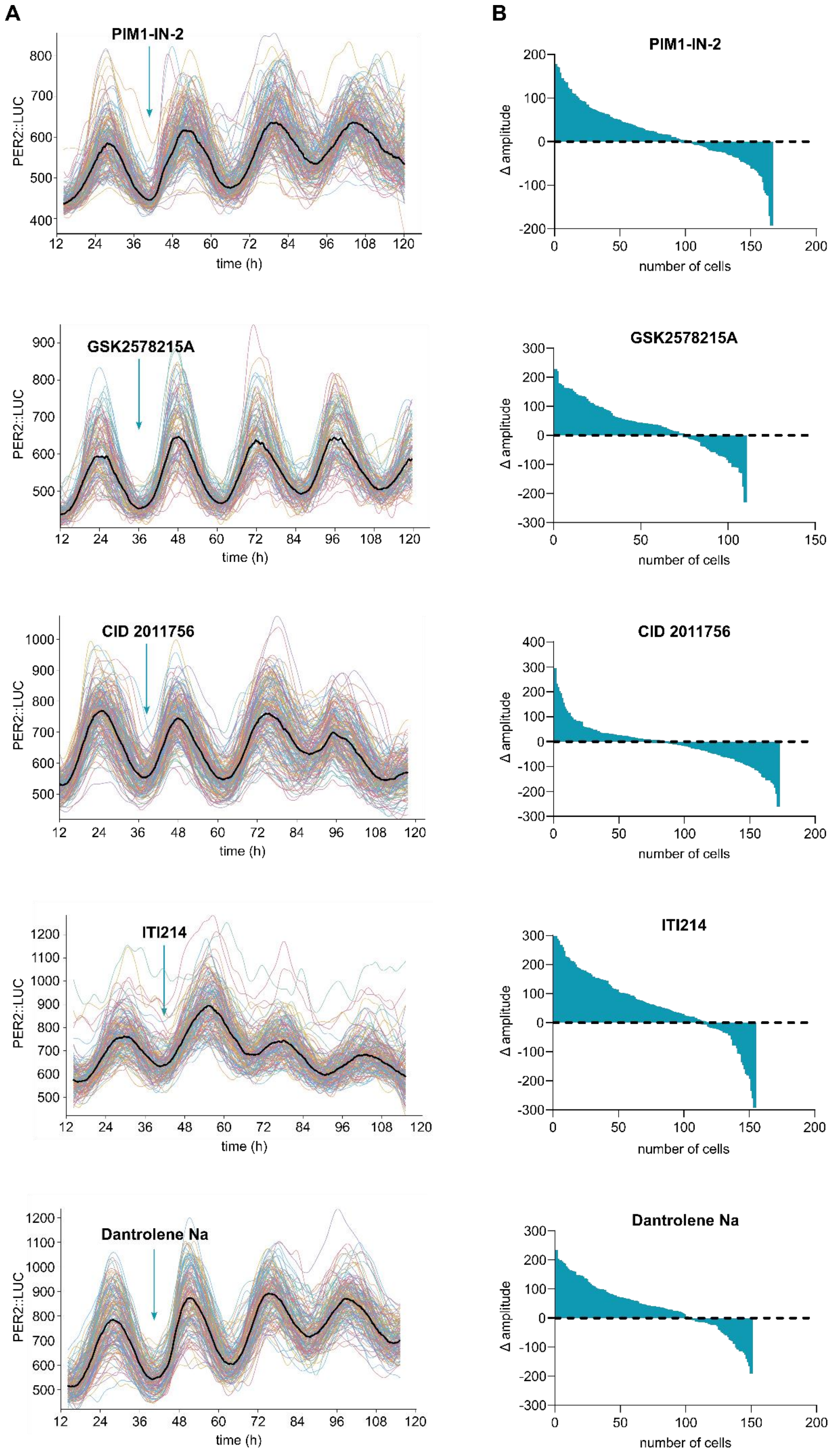
A
B
PIM1-IN-2
PER2::LUC
time (h)
Δ amplitude
number of cells
GSK2578215A
CID 2011756
ITI214
Dantrolene Na

**Fig. S5: Effect of hits on PER2::LUC rhythm amplitude in SCN-organotypic slices**

(A) Representative circadian rhythms of PER2::LUC expression in single cells from a suprachiasmatic nucleus (SCN) *ex vivo* explant obtained from a mouse brain, upon the application of hit compounds: PIM1-IN-2, GSK2578215A, CID 2011756, ITI214, or Dantrolene Na at 30 μM. Individual light traces: single cell rhythms, solid black line: moving average of all traces from a slice.

(B) Quantification of changes in the amplitude of PER2 rhythm in cells (correspondence to A). The difference in amplitude between before and after treatment with PIM1-IN-2, GSK2578215A, CID 2011756, ITI214, or Dantrolene Na is plotted against cell number, sorted by the magnitude of induced change in amplitude upon compound treatment. The point of intersection of the graph with the dashed line at Δ amplitude = 0 marks the transition from cells showing enhanced amplitude to cells with decreased amplitude of their rhythms after treatment.

### SI: Molecular structures of identified hit compounds

**VER-49009**

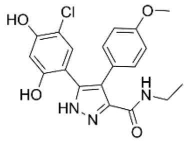

**OSI-420**

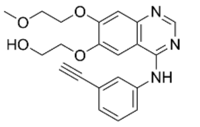

**Flutamide**

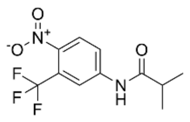

**SGC-CBP 30**

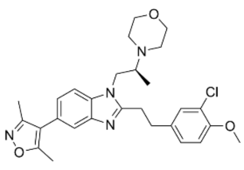

**EED226**

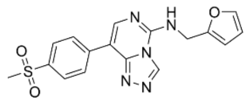

**WHI-P154**

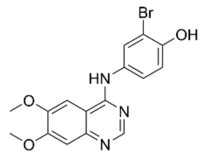

**ITI-214**

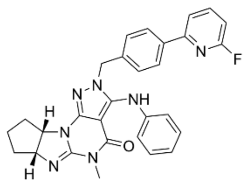

**PIM1-IN-2**

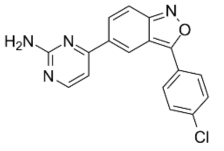

**CID 2011756**

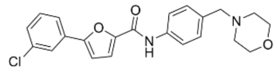

**Dantrolene Na**

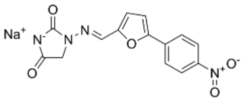

**Ulipristal acetate**

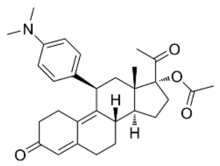

**Diclazuril**

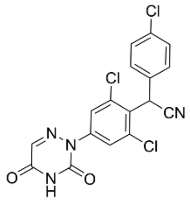

**GSK2578215A**

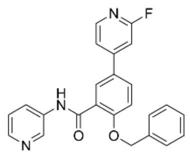

**Table S1: Amplitude and Period changes induced by hit compounds in NIH3T3 cells**

| Compound | % Amplitude enhancement above DMSO | | | Period (h) | | |
|---|---|---|---|---|---|---|
| | ***0.5 µM*** | ***1.5 µM*** | ***5 µM*** | ***0.5 µM*** | ***1.5 µM*** | ***5 µM*** |
| **DMSO** | | | | 25.4±0.72 | | |
| **CEM3** | 44.6±11.3 | 50.3±5 | 68.5±4.1 | 25.2±0.14 | 25.4±0.06 | 25.3±0.37 |
| **Isrib Transisomer** | 10.6±17.5 | 11.1±25.9 | 15.7±22.1 | 25.6±0.23 | 25.7±0.2 | 25.5±0.23 |
| **ITI-214** | 24.1±17.2 | 60±13.2 | 110±5.7 | 25.5±0.24 | 25.3±0.15 | 25.1±0.35 |
| **WHI-P154** | 29.5±11.9 | 38.4±19.2 | 50.6±17.8 | 25.4±0.21 | 25.3±0.12 | 25.1±0.07 |
| **VER49009** | 33.7±26.9 | 83.1±9.25 | 213±12.4 | 25.3±0.34 | 24.8±0.33 | 24.9±0.04 |
| **Flutamide** | 46.6±21.6 | 45.7±8.51 | 67.4±20.3 | 25.6±0.22 | 25.4±0.18 | 25.4±0.31 |
| **OSI-420** | 40±13.7 | 66.9±12.6 | 108±4.4 | 25.6±0.33 | 25.3±0.07 | 25.1±0.03 |
| **Ulipristal Acetate** | 13.8±16.9 | 22.8±21.7 | 32.2±15.5 | 25.5±0.3 | 25.4±0.29 | 25.4±0.21 |
| **PD53035** | -9.9±20.1 | -16.1±12.6 | 17.3±8.1 | 25.7±0.57 | 25.7±0.44 | 25.7±0.26 |
| **PIM1-IN-2** | 31.9±23.8 | 36.5±14.8 | 43.7±15.5 | 25.6±0.05 | 25.8±0.09 | 25.9±0.27 |
| **PQ401** | -11±5.2 | -5.1±2.6 | -2.8±3.8 | 26.2±0.11 | 25.9±0.03 | 26±0.16 |
| **Ruxolitinib** | 7.4±7 | 22.6±9 | 20±10.1 | 25.9±0.17 | 25.9±0.01 | 26±0.12 |
| **Dantrolene Na** | 12.8±11.9 | 35.2±13.6 | 52.5±7.8 | 25.4±0.22 | 25.4±0.11 | 25.4±0.19 |
| **Bay K 8644** | 6.3±7 | 23±16.4 | 23.3±2.9 | 25.2±0.19 | 25.2±0.08 | 25.3±0.11 |
| **CID 2011756** | 4.5±27.2 | 21.9±6.4 | 53.7±11.9 | 25.1±0.22 | 25.4±0.27 | 25.3±0.19 |
| **GSK2578215A** | 8.67±19.6 | 26.2±10.6 | 31.1±13 | 25.1±0.55 | 25.3±0.36 | 25.3±0.33 |
| **Diclazuril** | 28.3±18.3 | 38.6±9.3 | 42.3±12.5 | 25.1±0.34 | 24.9±0.19 | 25±0.2 |
| **SGC-CBP 30** | 11.9±18.8 | 43.8±5.6 | 41.4±6.6 | 25.5±0.2 | 25.4±0.2 | 25.8±0.08 |
| **ON123300** | 2.5±7.1 | -3.5±4.9 | -19.2±7 | 25.1±0.36 | 25.2±0.29 | 25.5±0.41 |
| **EED226** | 25±11.2 | 27.9±5.8 | 27.2±17.9 | 25±0.14 | 25.2±0.15 | 25.6±0.11 |

**Movie S1: Bioluminescence recording of PER2::LUC rhythm in SCN-organotypic slices upon treatment with WHI-P154**

Compilation of bioluminescence images (imaged every 30 mins) showing PER2::LUC circadian rhythms from a suprachiasmatic nucleus (SCN) *ex vivo* explant obtained from a mouse brain, upon the application of hit compounds WHI-P154 at 30 μM.
